# Library size can undermine accurate molecular and phenotypic subtyping in spatial transcriptomics data

**DOI:** 10.1101/2024.07.07.602370

**Authors:** Natalie C Fisher, Sudhir B Malla, Nigel B Jamieson, Philip D Dunne

**Affiliations:** The Patrick G Johnston Centre for Cancer Research, Queen’s University Belfast, Belfast, United Kingdom; Cancer Research UK Scotland Institute, Glasgow, United Kingdom; Wolfson Wohl Cancer Research Centre, Institute of Cancer Sciences, University of Glasgow, Glasgow, United Kingdom

## Abstract

In an era where transcriptomics-based subtyping, phenotyping and mechanistic understanding is increasingly being driven by state-of-the-art spatially resolved transcriptomic (ST) technologies, it is imperative that researchers, journals, and funders do all they can to ensure that as a community we are interpreting these exciting data as accurately as possible with awareness of their limitations. In this short report, we highlight one potential bias in ST data that could undermine accurate <u>interpretation</u> of transcriptional signatures, providing the field with an opportunity to identify and avoid this issue prior to release of new mechanistic findings. This issue is particularly relevant for platforms that produce some of the most granular and high-resolution spatial information at single cell (and sub-cellular) resolution, with the compromise of a reduced transcriptome panel of genes (**Figure 1A**).

## Background

As ST technologies are still in the early competition stages of development, subject to rapid release and obsolescence, the range of available platforms will continue to evolve^***1***^ before reaching a stabilised state. Alongside this expansion, benchmarking studies are objectively assessing the range and dynamics of gene expression across platforms^2-4^ with their performances/stats being comprehensively reviewed^5^. As recently described, its particularly important to process spatial data in a way that considers library size.^6^ To complement the need for specific normalisation protocols due to reduced library size, we believe that the field must also ensure that the biological interpretation of existing transcriptional signatures/subtypes in the same reduced gene panels remains faithful to the same mechanistic signalling or clinical features they represent in non-spatial data.

This is particularly important at this point in time, as ST data offers the opportunity to uncover even more novel and “*seen-for-the-first-time*” mechanistic insight afforded by the complex spatial data points at multiple resolution scales; including cellular neighbourhoods, single-cells, subcellular RNA localisation. Any deviation in mechanistic meaning of even the most well-validated and routinely used signatures/subtypes at this stage will therefore lead to exponential levels of misunderstanding.

### Potential issue with interpreting existing gene signatures within ST studies

While ST platforms will continue to expand the gene panels they offer, mechanistic biologists that seek single cell resolution must accept the compromise that anywhere up to 95% of the transcriptome may not be represented on current ST panels (**Figure 1B**), an example of this is the ∼1000 gene panel available on the Nanostring CosMx platform. Although existing transcriptional signature/subtype scores, developed from full transcriptome bulk and/or single cell data, can still be enumerated in individual cells from a ST dataset, the ability to interpret these signatures scores in the same way in a reduced transcriptome panel compared to what they would be in a whole transcriptome dataset, remains understudied.

**Figure 1.**
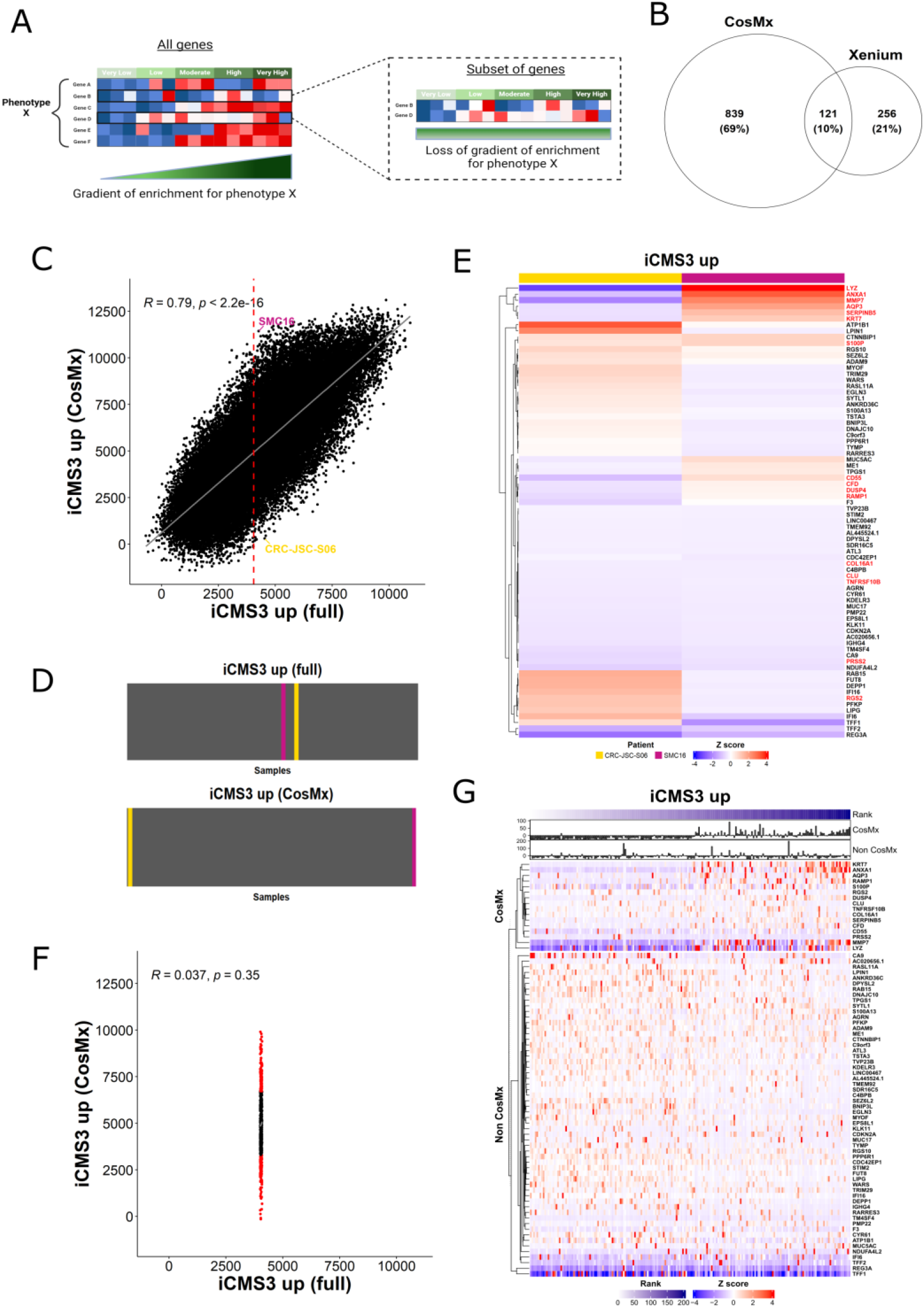
Visualisation of the iCMS3 up signature. A: Schematic overview of enrichment using a full gene panel versus a reduced panel. B: Overlap between the genes represented on the CosMx and Xenium gene panels. C: Correlation between the single sample scores of the full iCMS3 up signature (74 genes) and the 15 genes from the signature represented on the CosMx platform. Two samples are highlighted which have a similar iCMS3 up enrichment for the full signature (CRC-JSC-S06: 4535.746; SMC16: 4265.263) but extreme enrichments for the signature composed only of the genes present on the CosMx array (CRC-JSC-S06: 301.0396; SMC16: 11430.593). Median (4047.994) shown by red line. D: Visualisation of the rank of each sample across for the full (CRC-JSC-S06: position 25872/44458; SMC16: position 23907/44458) and CosMx (CRC-JSC-S06: position 513/44458; SMC16: position 44241/44458) signature enrichment. E: Heatmap showing the relative enrichment of each gene with the iCMS3 up signature for each sample, with the genes present on the CosMx array in red. F: Subset of samples (n=628) +/-1% of the median (4007.514 – 4088.474), with the top 100 and bottom 100 samples for CosMx enrichment in red. G: Heatmap of the top 100 and bottom 100 samples (shown in red in F) for CosMx enrichment with a full iCMS3 up enrichment around the median. The samples are arranged by the rank of each sample for CosMx signature enrichment, with the sum of the genes within the iCMS3 up signature present on the array (CosMx [n=16]) and those not present on the array (Non CosMx [n=58]) overlaid as barplots.

### Examples using existing data

To test concordance of outputs that would be obtained using a condensed ST panel compared to a wider full transcriptome panel, we performed an assessment of single epithelial cell transcriptional scores from an independent scRNAseq cohort. Using these data, we performed a comparison of outputs from the full (reference) signature scores to those derived when using only the genes on a condensed ST panel. The example below offers a typical situation, where single cell scoring with one of the iCMS transcriptional signatures^7^ is performed using whole transcriptome data (labelled ‘full’) and when filtered to only include genes currently offered on the standard CosMx 1000 gene panel (labelled ‘CosMx’).

Comparing the iCMS3 up signature scores derived from the full transcriptome (which includes all n=74 iCMS3 up genes) to the score that would be derived using only the genes present on the CosMx reduced panel (which includes only n=16 iCMS3 up genes), reveals an overall general correlation of the scores (R = 0.79; p < 2.2×10^−16^). Importantly however, despite population-wide correlation, given the spread of the data there are numerous cases where cells switch from being identical scores in one technology, to remarkably extreme opposites in the genes provided in the other technology. In a clear example of this issue, we selected pairs of cells that provide remarkably similar iCMS3 scores when using the full iCMS3 up signature, as they will have been described in the original iCMS study, yet when assessed using the reduced gene panel available in the CosMx, these cells would be classified as having the most extreme opposite top/bottom scores.

This can lead researchers to interpret these data in entirely opposing directions to the originally intended mechanistic meaning, as outputs for these cells would be classified as either having almost <u>exactly the same</u> biological phenotype if profiled using a full transcriptome panel, or the <u>most extreme and entirely opposing biologies</u> if profiled using the 1000 gene CosMx panel (**Figure 1C, 1D**).

### What is the reason for this

This discordance arises as a signature/subtype score summarises all gene expression values within that specific signature, and while these two cells/samples have the same iCMS3 up full transcriptome signature score, it is via elevation of entirely different combinations of signature genes, as revealed by their unique patterns of gene expression and extreme signature enrichment rank in the genes present on the CosMx platform (1000 gene panel) coloured in red (**Figure 1E**). When this alternating pattern of individual gene expression overlaps with the genes that are represented/excluded on the CosMx panel, as above, the outputs are severely altered.

Furthermore, when we select all cells with a full transcriptome iCMS3 up score around the population median (4047.994 +/-1% expression units, n=628 cells), although these cells all have similar full iCMS3 scores, when we assessed their complementary CosMx iCMS3 up scores in parallel, they ranged across the full spectrum, from the highest 4.9% to the lowest 0.8% of all cells in this cohort (**Figure 1F**; lowest score: 149.02554 [375/44458 samples with a score ≤ 149.02554]), (highest score: 9905.982 [2194/44458 samples with a score ≥ 9905.982]).

From these, we next selected the top and bottom n=100 subsets of these cells ranked by CosMx score (**Figure 1F**) and examined in detail the individual gene expression values of the iCMS3 signature genes. These data were stratified according to whether they were represented solely within the full gene list (non-CosMx) or also on the CosMx panel, which reveal that while these cells/samples all would be classified as having the same iCMS3 up score in a scRNAseq experiment, the expression patterns of the genes on the CosMx panel are highly distinct, leading to a discordant CosMx result (**Figure 1G**).

### Bias in this current study

By using the same source data, altering only the number of genes included, we have selected favourable conditions for identifying similarities across these two axes. Therefore, our study will almost certainly <u>underestimate</u> discordance, as it avoids well-documented differences between sequencing and normalisation methods particularly in reduced library size panels.^6^ To highlight the more extreme effect of applying signatures which have been developed independently of the discovery data, we conducted the same analyses in the MYC and PRC signatures^8^, which form the basis of our recently published PDS molecular subtypes.^9^ Despite a strong overall correlation (**Figure 2A,C**) samples with a median full signature again span almost the entire range of CosMx scores (**Figure 2B,D**). The extreme effect of the specific subset of genes present on the CosMx array has on the enrichment score compared to considering the full signature is also illustrated using the MYC and PRC signatures (**Supplementary Figure 1**). In addition, single cell data will be biased towards exclusion of necrotic or broken/ruptured cells^10^ which may remain present in spatial data. Therefore, there is likely to be much more variability between subtype/signature interpretation from ST and non-ST data than documented here. While this current situation may seem like it is championing the use of wider panel region-of-origin methods, like 10X Genomics VISIUM or Nanostring GeoMx, such approaches must themselves continue to ensure that interpretation of signature scores are viewed based on confounding lineages^11^ or biologically heterogeneous pools of phenotypically distinct lineages^9,12^ compounded by biased region of interest selection strategies.^13^

**Figure 2.**
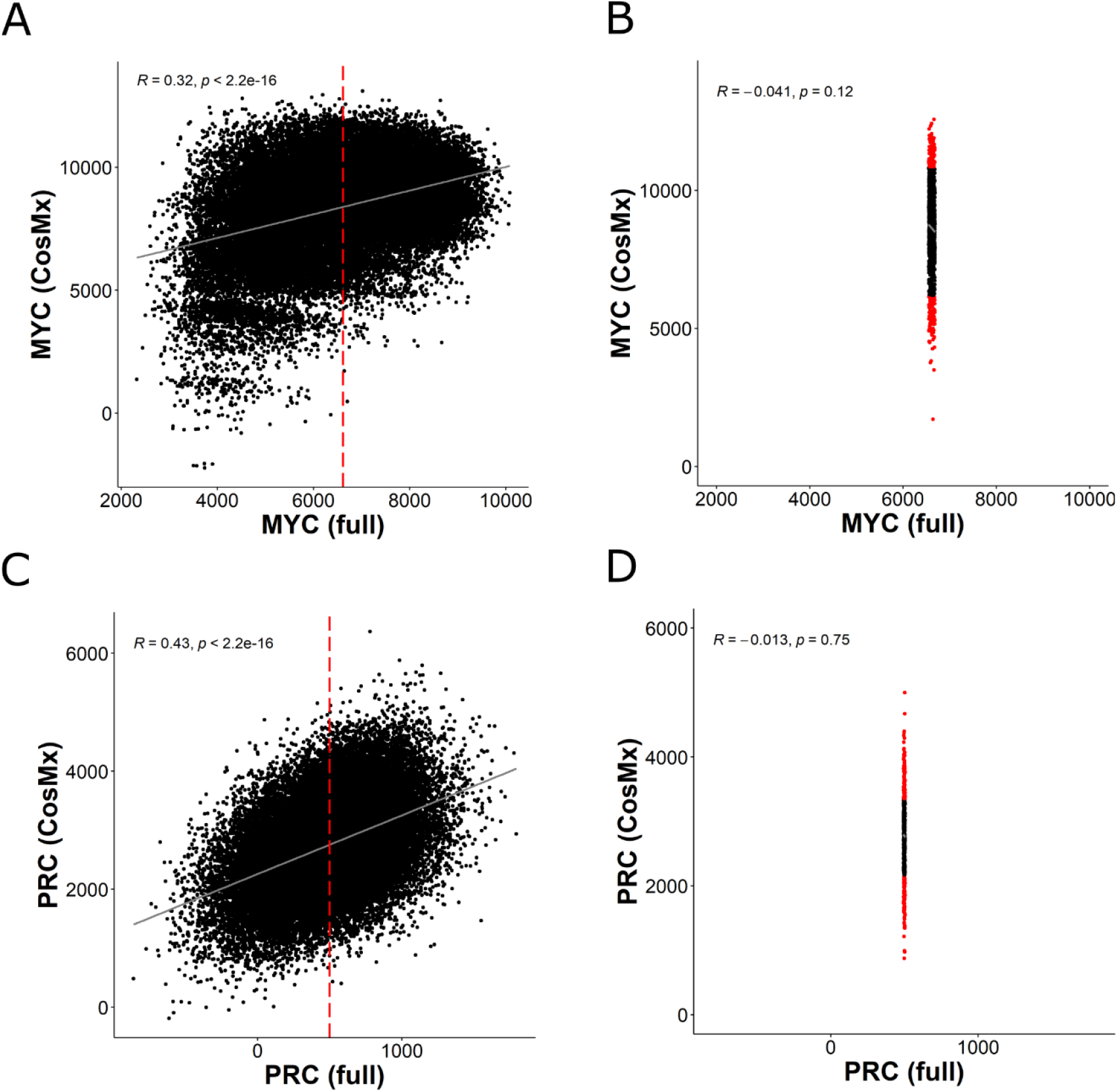
Visualisation of the signatures underlying PDS biology. A: Correlation between the single sample enrichment of the full MYC (291 genes) and the 9 genes from the signature represented on the CosMx platform. Median (6616.096) marked by the red line. B: Subset of samples (n=1426) +/-1% of the median (6549.935 - 6682.257), with the top 100 and bottom 100 samples for CosMx MYC enrichment in red. C: Correlation between the single sample enrichment of the full PRC (395 genes) and the 42 genes from the signature represented on the CosMx platform. Median (498.9944) marked by the red line. B: Subset of samples (n=591) +/-1% of the median (494.0045 – 503.9843), with the top 100 and bottom 100 samples for CosMx PRC enrichment in red.

### Moving forward

We reside within an era where molecular information from even the most advanced profiling technologies is easier and cheaper to generate than ever before. As such, new insights into cancer mechanisms that examine individual genes or multiple signalling pathways at an incredibly high resolution are routinely being published, which, with informed interpretation, can reveal new understanding of the molecular events underpinning tumour initiation, progression, and response to treatment.

As we move towards imaging-based transcriptomic assessment, a trade off exists between gene panel number and tissue imaging time. The current coverage of genes for standard panels will no doubt continue to increase, but this sits at ∼400 genes for single cell-specific ST approaches (Xenium [10x Genomics]) and currently ∼1000 genes (CosMx [Nanostring]) as standard (∼6000 genes recently launched), with up to ∼18,000 genes for the regions-of-interest ST options (GeoMx [Nanostring]) or unbiased whole transcriptome (Visium, Visium HD [10x Genomics]. Importantly however, with these latter approaches you lose specificity of the cell type being profiled or the heterogeneity in regions compared to cell-specific technologies. Although large 18,000 gene panels at a single-cell spatially orientated resolution are on the horizon, as it stands, ST data either provides wide transcriptome in non-specific cells, or more reduced transcriptomes in specific cells. Until there is full transcriptomic coverage, data presented here indicates that existing signature/subtype results derived from current reduced transcriptome ST panels should not be immediately interpreted as providing the same mechanistic understanding of signatures/subtypes from other benchmarked methods.

We believe this issue may go unnoticed in some studies, as many of the most commonly used algorithms/tools will still produce a signature/subtype score regardless of how many of the template genes are present, as they simply “ignore” genes from a specific signature that are not present in a dataset^14^. There are remarkable advantages of using signatures/subtypes and pathway scores to represent intricate multi-gene cascades, particularly coupled with the resolution of ST data. However, as demonstrated above, in their current form they may not always represent the same phenotypic cascades in ST data. In the interim we have developed SpatialExploreR, (https://mallsid.shinyapps.io/spatialexplorer/), an online resource that enables users to assess the impact the reduced gene panel (CosMx) has on the enrichment of a suite of commonly used transcriptional signatures and subtypes. This resource can be used via any web browser, to interrogate the data and signatures we have described here, however, as we have also released the code underpinning the resource, users can embed any other datasets and/or signatures relevant to their own field for similar testing.

In an attempt to find a surrogate for signature enrichment in the limited CosMx gene panel we correlated the enrichment score per sample with all genes represented on the panel (n=960), but neither the single most correlated gene, nor the sum of the top five most correlated genes improve upon the correlation observed with the reduced signature enrichment (**Supplementary Figure 2**). Therefore our findings strongly support the development of a new wave of experimentally validated signatures designed specifically for spatial phenotypic characterisation.

Overall, the field is at a crossroads, where generation of high-resolution data continues rapidly and at scale, however the intricacy of the molecular information and the diverse analytical methods required have created a critical bottleneck for non-programmers to interpret these data,^13^ which now demands specialised computational skills and expertise. Although funders have invested significant resources in generating large-scale molecular datasets, the value and impact of these resources are limited if only a small subset of researchers and clinicians have the necessary computational skills to analyse and interpret the data effectively. Given the complexity of these technologies, their widespread use must also be coupled with commitments from funders for substantially more support for data-driven early career researchers (ECRs), as these ECRs are the ones who have the skills and expertise to analyse the data effectively and most importantly to interpret these findings accurately.

## Methods

### Data Processing

The processed count expression matrix for epithelial cells and corresponding epithelial metadata were downloaded through the Synapse under accession code syn26844071^7^. The count expression matrix for tumour epithelial cells (n = 44458) was normalized using *SCTransfrom* in the *Seurat* R package^15^ (v.4.3.0.1) with variable.features.n = 26078 to account for all genes.

### Gene lists

The iCMS genesets were downloaded from the original publication^7^. The MYC and PRC signatures detailed with the PDS molecular classification^16^ were downloaded from the original source^8^. A signature collection was created for all genes within the signature (full signature) and those which overlapped with genes (n = 960; default Nanostring exprMat file) on the CosMx gene panel (CosMx signature).

### Single Sample Geneset Enrichment

Single sample GSEA scores were calculated per cell for each signature using the *enrichIt* function in the *escape* R package ^17^(v1.8.0) with groups = 1000, cores = 2, and min.size=5.

### Correlation

The gene expression matrix was filtered to only include the genes represented on the CosMx panel. The correlation between the MYC, PRC and iCMS up ssGSEA scores for the full signature were calculated for each gene using the *cor*.*test* function with method = pearson in the *stats* R package (4.2.3).

### Data visualisation

Data visualisations was processed using R (v4.2.3) in RStudio. Packages used included *VennDiagram* (v1.7.3), *ggpubr* (v0.6.0), *ggplot2* (v3.5.0), *circlize* (v0.4.15) and *ComplexHeatmap* (v2.14.0). Pearson’s correlation was calculated with in the *ggpubr* (v0.6.0) R package.

### SpatialExploreR shiny application

The shiny-based web application was built on using list of shinyapps related R packages, including *shiny* (v1.8.1.1), *shinydashboard* (v0.7.2), *shinydashboardPlus* (v2.0.4), *shinyWidgets* (v0.8.6), *shinyjs* (v2.1.0) and *shinycssloaders* (v1.0.0). Other R packages for data visualisation includes, *ggplot2* (v3.5.1), *plotly* (v4.10.4), *ComplexHeatmap* (v2.18.0), and *InteractiveComplexHeatmap* (v1.10.0).

### Code for all results in GitHub

All the scripts, including for the ShinyApp will be deposited during review process.

## Supplementary Figures

**Supplementary Figure 1.**
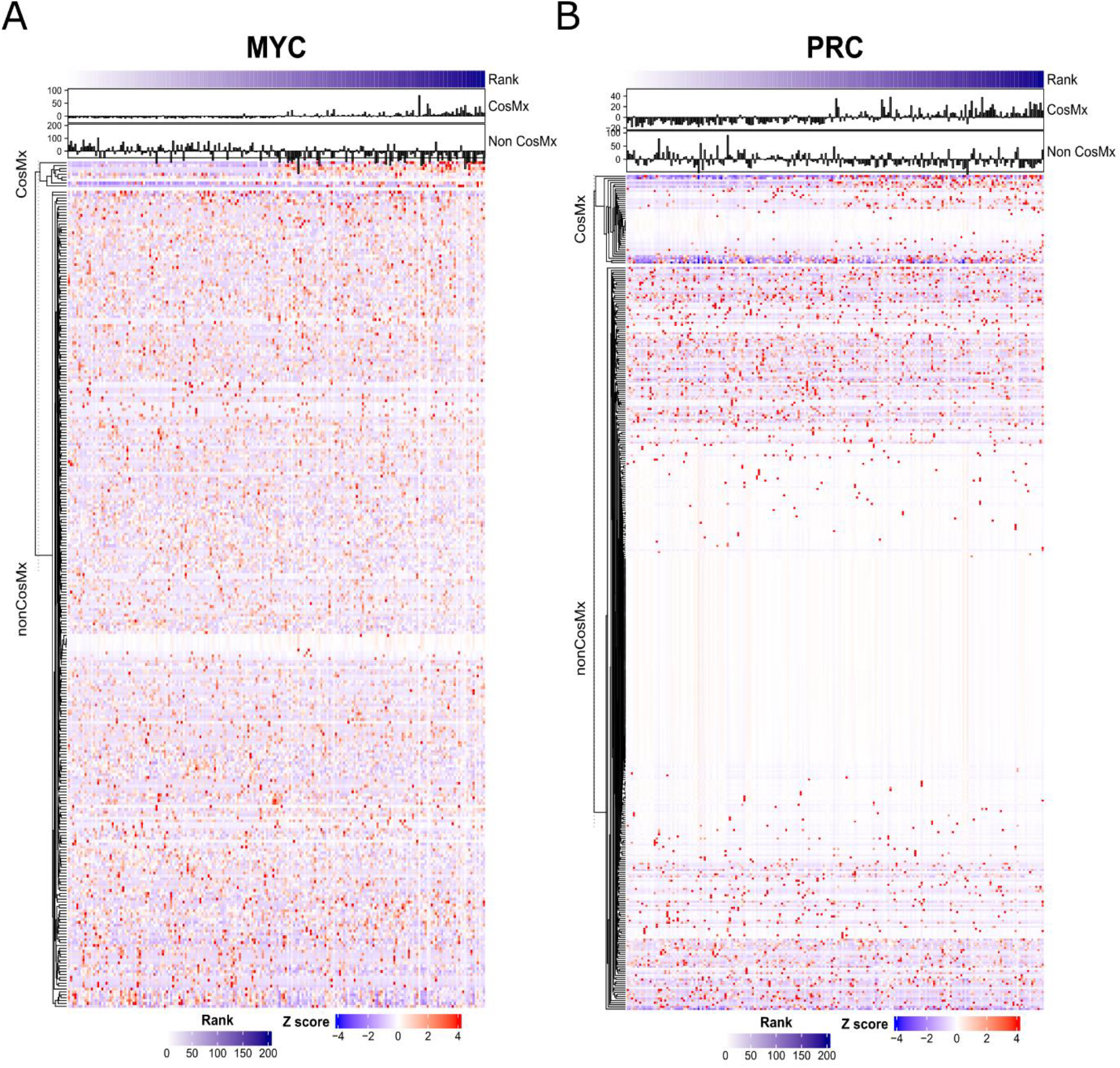
Contribution of genes within MYC and PRC signatures to overall sample enrichment. A: Heatmap of the top 100 and bottom 100 samples (shown in red in Figure 3B) for CosMx enrichment with a full MYC enrichment around the median. The samples are arranged by the rank of each sample for CosMx signature enrichment, with the sum of the genes within the MYC signature present on the array (CosMx [n=282]) and those not present on the array (Non CosMx [n=9]) overlaid as barplots. B: Heatmap of the top 100 and bottom 100 samples (shown in red in Figure 3D) for CosMx enrichment with a full PRC enrichment around the median. The samples are arranged by the rank of each sample for CosMx signature enrichment, with the sum of the genes within the PRC signature present on the array (CosMx [n=353]) and those not present on the array (Non CosMx [n=42]) overlaid as barplots.

**Supplementary Figure 2:**
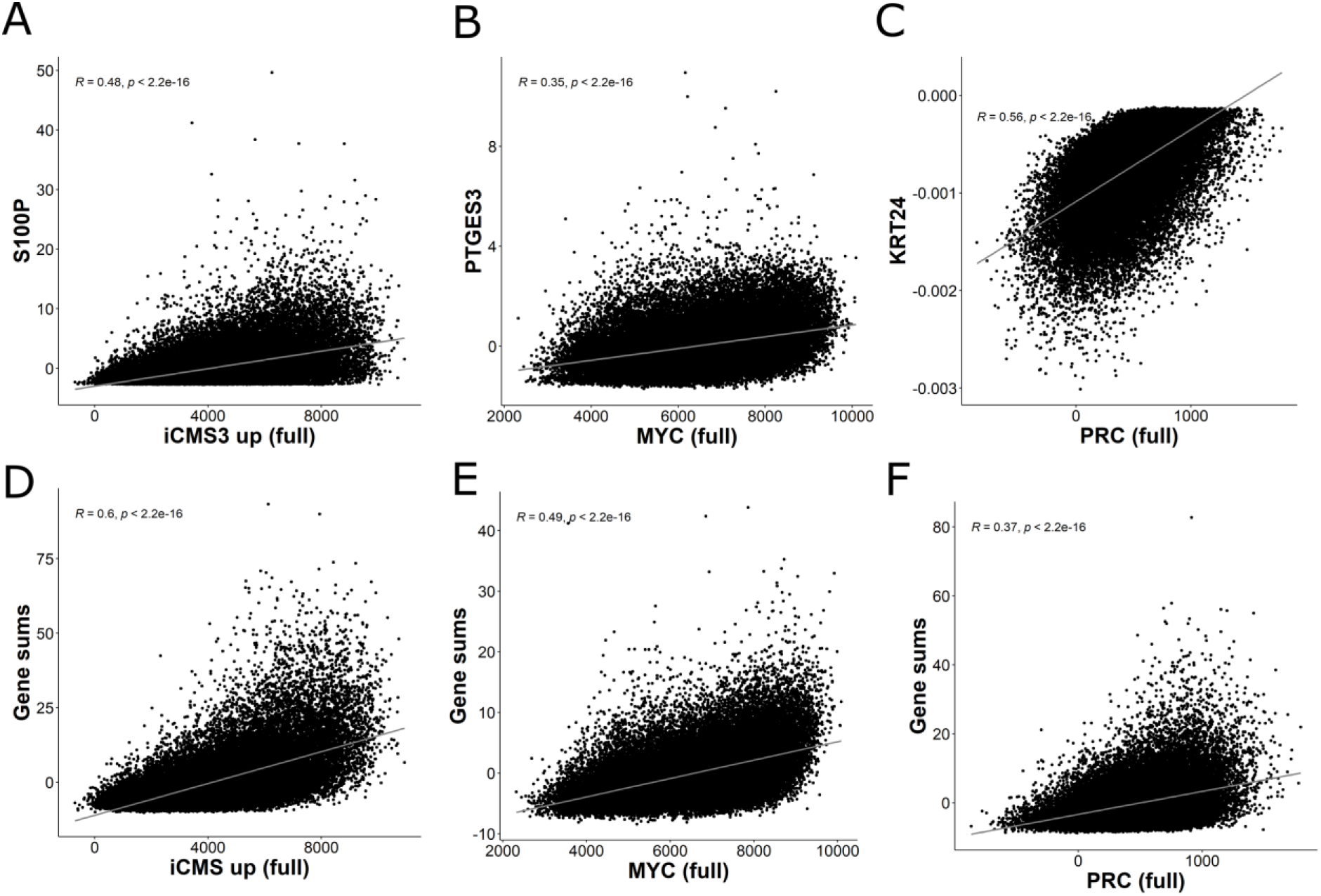
Correlation between signature enrichment and genes on CosMx. The most correlated gene (A-C) and the sum of the top five most correlated genes (D-F) are correlated against the single sample enrichment score of the full iCMS up (A,D), MYC (B,E) and PRC (C,F).

