## Supplementary figures and images for "Library size can undermine accurate molecular and phenotypic subtyping in spatial transcriptomics data"

### Supplementary Figure 1

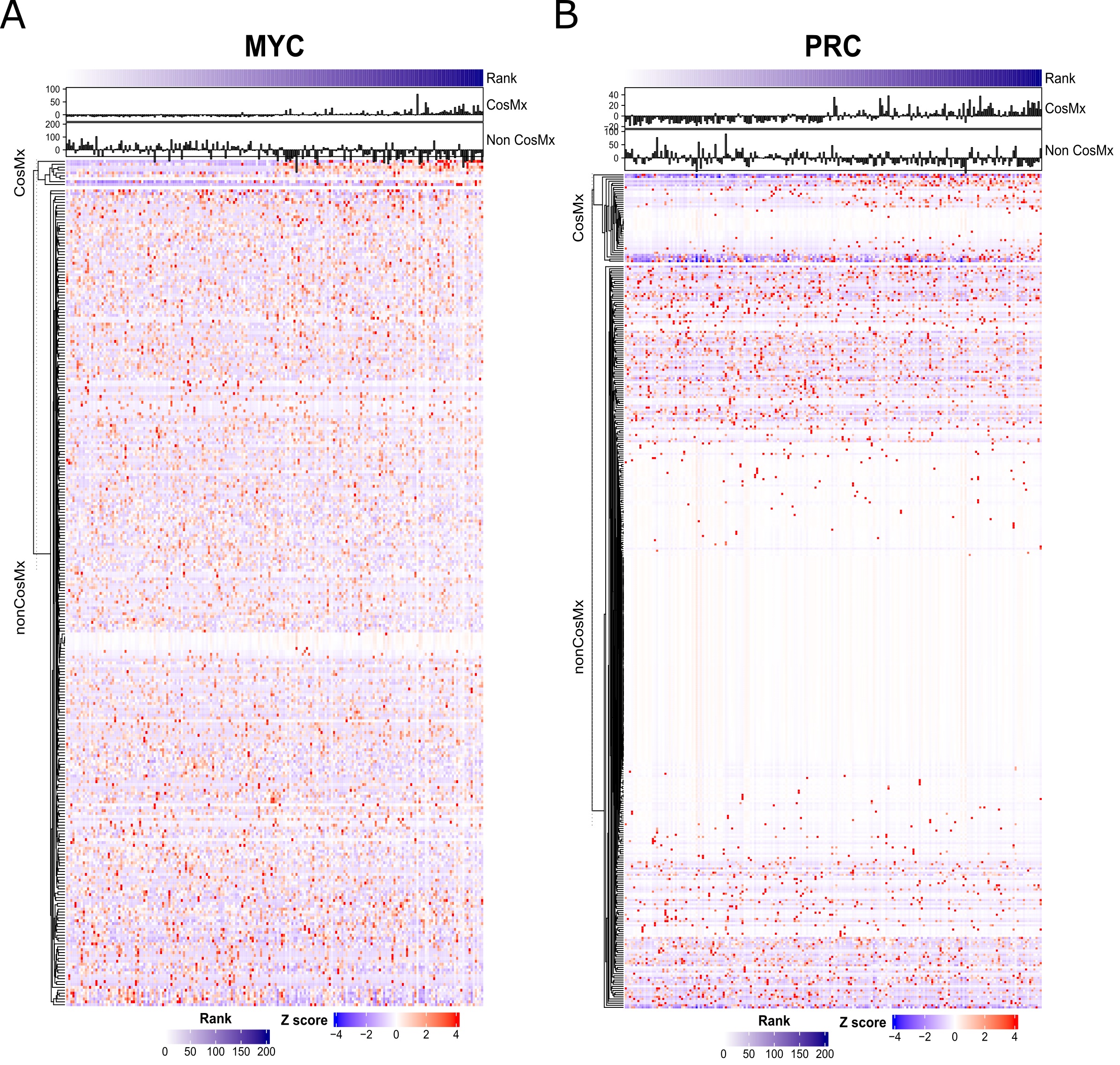

### Supplementary Figure 2

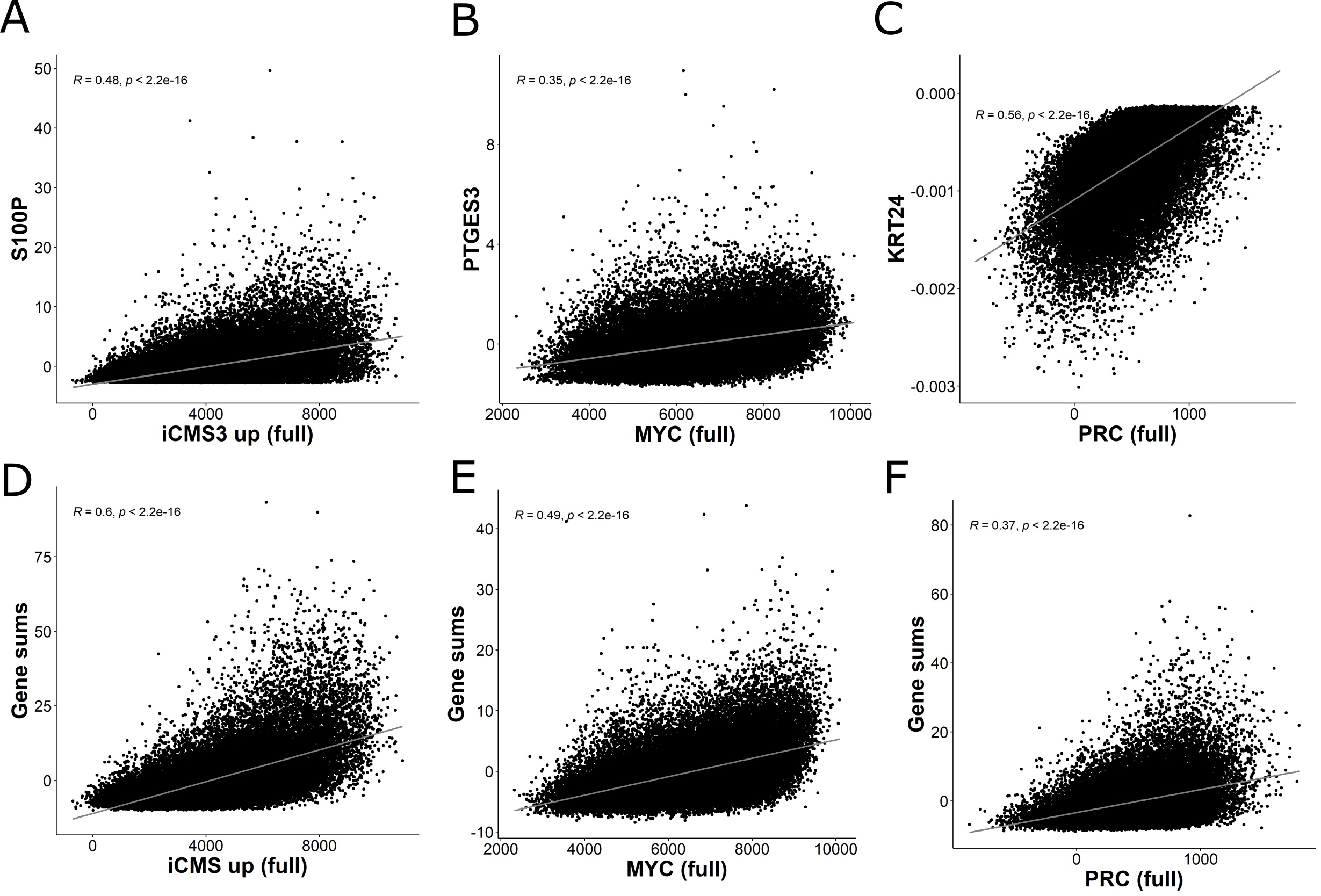
